# Effects of muscarinic M_1_ receptor stimulation on reinforcing and neurochemical effects of cocaine in rats

**DOI:** 10.1101/2020.03.10.972687

**Authors:** Pia Weikop, Kathrine L. Jensen, Morgane Thomsen

## Abstract

Cocaine addiction is a chronic illness characterized by maladaptive drug-induced neuroplastic changes that confer lasting vulnerability to relapse. Over several weeks we observed the effects of the M_1_ receptor-selective agonist VU0364572 in adult male rats that self-administer cocaine in a cocaine vs. natural reinforcer choice procedure. The drug showed unusual long-lasting effects, as rats gradually stopped self-administering cocaine, reallocating behavior towards the food reinforcer. The effect lasted as long as tested and at least four weeks. To begin to elucidate how VU0364572 modulates cocaine self-administration, we then examined its long-term effects using dual-probe in vivo dopamine and glutamate microdialysis in nucleus accumbens and medial prefrontal cortex, and ex vivo striatal dopamine reuptake. Microdialysis revealed dramatic decreases in cocaine-induced dopamine and glutamate outflow four weeks after VU0364572 treatment, without significant changes in dopamine uptake function. These lasting and dramatic effects of M_1_ receptor stimulation reinforce our interest in this target as potential treatment of cocaine addiction. M_1_ receptors are known to modulate medium spiny neuron responses to corticostriatal glutamatergic signaling acutely, and we hypothesize that VU0364572 may oppose the addiction-related effects of cocaine by causing lasting changes in this system.

## INTRODUCTION

Drug addiction is a chronic relapsing illness characterized by complex changes in the biochemistry of the brain, many of which persist after weeks or months of abstinence and confer lasting susceptibility to relapse [1,2]. Signaling in the prefrontal cortex (PFC), and from the PFC to the nucleus accumbens (NAc), and downstream dopamine signaling in NAc medium spiny neurons (MSNs), is considered a main pathway of the addicting effects of abused drugs including cocaine [3]. Central to this “addiction pathway” are glutamatergic corticostriatal projections from PFC to NAc, which become dysregulated after chronic drug use, resulting in loss of control over drug taking behaviors [3–6]. The complex long-term adaptations induced by cocaine are not fully understood, but clinical and preclinical studies indicate that basal PFC activity and glutamate levels are reduced while responses to cocaine and cocaine cues are exaggerated [7,8]. Less scrutinized in addictions is the muscarinic cholinergic system. Yet activity of cholinergic interneurons, which provide the major acetylcholine input in striatum/NAc, is related to stimuli that predict reward / reward omission, and indeed is more responsive to changes in reinforcement contingency than the mesolimbic dopaminergic neurons, whose reward prediction error encoding is well described [9–11]. Muscarinic acetylcholine M_1_ receptors are the most abundant muscarinic receptor in neocortex and striatum and are expressed, mainly postsynaptically, on most if not all neocortical pyramidal neurons and striatal MSNs including MSNs in NAc [12–17]. Activity at M_1_ receptors modulates corticostriatal glutamatergic signaling, integration of inputs by MSNs, and neuroplasticity, and appears to preferentially facilitate inhibition of behavior [18–21]. M_1_ receptors are thus well placed to modulate some of the processes affected by cocaine exposure.

We have previously shown that acute pharmacological stimulation of M_1_ receptors can fully suppress cocaine self-administration behavior in mice, while food-maintained behavior was unaffected [22]. The M_1_/M_4_ receptor-preferring agonist xanomeline reduced cocaine taking in rats given access to choose between cocaine or a palatable food reinforcer [23]. Unlike effects of dopamine receptor ligands in the same assay [24,25], the effects of M_1_/M_4_ stimulation grew larger during subchronic dosing, and subsisted briefly after ended treatment [23]. In mice, M_1_/M_4_ stimulation facilitated extinction of cocaine seeking and prevented reinstatement of cocaine seeking when administered during extinction or between cocaine exposure and extinction [26]. Those findings suggested an unusual and possibly lasting modulation of the addiction-related effects of cocaine by M_1_/M_4_ stimulation, but the mechanisms are largely unknown. Here, we sought to delineate and better understand the contributions of M_1_ receptors to these effects. We assessed how the highly selective, brain-penetrant M_1_ receptor agonist VU0364572 [27,28] affected cocaine self-administration behavior over time in rats, using the cocaine vs. food choice assay. To begin to understand how VU0364572 modulates cocaine self-administration, we then examined its long-term effects using in vivo dopamine and glutamate microdialysis in NAc and medial PFC (mPFC), and ex vivo striatal dopamine reuptake.

## MATERIALS AND METHODS

### Animals

Behavioral studies were conducted at McLean Hospital in accordance with the NIH Guide for the Care and Use of Laboratory Animals and were approved by the McLean Hospital Institutional Animal Care and Use Committee, using experimentally naïve male Sprague Dawley rats acquired at age 7-8 weeks (Charles River, Wilmington, MA). Rats were group-housed up to 4/cage until catheter implantation, then housed individually. Water was available ad libitum, food was provided in amounts adjusted to maintain body weight at about 400-450g (≈17 g/day Rat Diet 5001; PMI Feeds, St. Louis, MO). For enrichment, species-appropriate treats were provided once or twice weekly. Microdialysis and dopamine uptake study procedures were conducted at the Laboratory of Neuropsychiatry in an AAALAC-accredited facility, in accordance with the EU directive 2010/63/EU and were approved by the Animal Experiments Inspectorate under the Danish Ministry of Food, Agriculture, and Fisheries. Experimentally naïve male Sprague Dawley rats aged 6 weeks were acquired from Taconic Denmark; they were pair-housed with ad libitum water and rodent chow (Altromin 1310, Brogaarden, Denmark). All rats were acclimated to the temperature- and humidity-controlled facilities for a week before experiments began. Lights were on from 07:00 to 19:00, testing occurred during the light phase. Rats were randomly assigned to experimental groups, and groups were evenly balanced between test stations and days/times. Whenever possible, the experimenter was blind to treatment group.

### Behavioral studies: cocaine self-administration

Rats were trained and tested in a cocaine vs. food choice procedure as previously described [23,29]. Rats were trained to self-administer intravenous cocaine by pressing one lever five times, or receive a liquid food treat by pressing another lever five times. At completed training, daily sessions consisted of five 20-min components during which cocaine dose increased for each component: 0, 0.056, 0.18, 0.56, 1.0 mg/kg/infusion, and food reward was held constant. Baseline behavior was recorded as the mean of three consecutive sessions, then, the effects of VU364572 were tested. First, acute effects of a 0.032 – 1.8 mg/kg were assessed, initially within-subjects with at least three sessions of baseline between doses. It became apparent that higher doses of VU364572 had lasting effects, and we then tested saline, 0.1 and 1 mg/kg VU0364572 in individual subjects, with up to four weeks observation periods after single dosing, during which cocaine and food reinforcer were available in daily sessions as during baseline conditions, five days/week (Fig.1A). The first rat treated with 1 mg/kg VU0364572 was only observed for two weeks, and three lost catheter patency during testing, resulting in a group size of n=6-9. Only data collected with demonstrated patent catheters were included.

**Figure 1.**
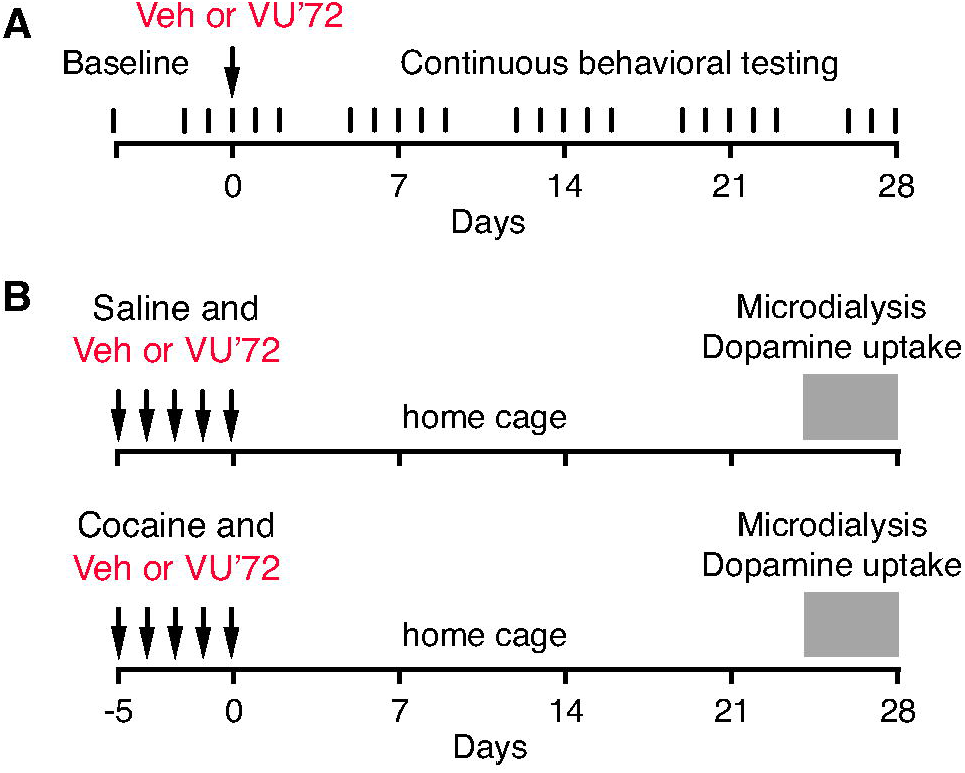
Experimental timelines. In the self-administration behavior experiment (A) rats were trained to baseline criteria over several weeks, then tested continuously 5 days per week for four weeks after administration of vehicle (Veh) or either 0.1 or 1 mg/kg of the M_1_ agonist VU0364572 (VU’72). For microdialysis and dopamine uptake studies (B), rats received saline (“cocaine-naïve” groups) or cocaine (“cocaine-experienced” groups), with vehicle or VU0364572.

### Microdialysis

Rats were injected once daily for five consecutive days with 1 mg/kgVU0364572 or vehicle, and, 30 min later, with 15 mg/kg cocaine or saline intraperitoneally, and left for one hour in a non home-cage enclosure. Treatment groups were: VU0364572+saline, VU0364572+cocaine, water+saline, water+cocaine (n=6). All rats were then left in the home cage to allow the effect VU0364572 to develop. Six age-matched controls, habituated to handling but otherwise naïve, were administered saline on the microdialysis day. For practical feasibility, rats were treated and tested in two cohorts on consecutive weeks, and microdialysis was performed over 5 days for each cohort. Microdialysis was thus performed 24-28 days after the last injection (Fig. 1B).

Surgery, microdialysis sampling, and neurotransmitter analysis were performed as previously described [30] with minor modifications. Two microdialysis probes (CMA/12 2mm, CMA/Microdialysis AB, Sweden) were implanted under sevoflurane anesthesia for simultaneous sampling of dopamine and glutamate in the mPFC (probe tip at AP, +2.5mm; ML, +0.5mm; DV, −5.4mm relative to Bregma) and NAc (AP, +1.8mm; ML, −1.4mm; DV, −7.4mm) in the same subjects. Anesthesia was maintained throughout testing. Probe placements were verified after ended experiments and are reported in Fig. S1 online. The first four 20-min samples after starting perfusion were discarded, then, three samples were used to determine baseline level, and five further samples were collected after intraperitoneal administration of 15 mg/kg cocaine. Relative concentrations of dopamine in the dialysate were determined as previously by HPLC with electrochemical detection [30]. Relative glutamate concentrations were detected with fluorescence HPLC, see Supplemental material online for full details.

### Dopamine uptake

A separate cohort of rats (n=4 per group, then repeated), was used for dopamine uptake, using the same treatment regimen as for microdialysis studies. Rats were sacrificed 24-27 days after the last injection (4 rats per day, counterbalanced by treatment group), and left ventral striatal tissue was dissected on ice from coronal sections using a brain matrix and 3mm punch. Synaptosomal dopamine uptake was assessed as previously described [31,32]. Background-subtracted fmol/min/μg protein counts were fitted by Michaelis–Menten kinetics using the median of triplicate determinations for each rat.

### Drugs

Cocaine hydrochloride was provided by the National Institute on Drug Abuse (behavioral studies) or purchased from the Copenhagen University Hospital pharmacy (Copenhagen, Denmark; Copenhagen studies). VU0364572 was synthesized as previously described at Vanderbilt University [28], and was generously provided by Drs. P. Jeffrey Conn and Craig Lindsley (Vanderbilt University). Cocaine was dissolved in 0.9% saline. VU0364572 was dissolved in sterile water, fresh daily, and administered intraperitoneally 30 min before sessions.

### Data analysis

Data were analyzed by analysis of variance with between- and within-subjects variables; for microdialysis, in addition, area under the curve (AUC) for 0-100 min after cocaine administration was calculated and analyzed. See supplemental materials for detailed descriptions and power analyses. Level of significance was set at α=0.05. Data are presented as group means ± standard error of the mean (s.e.m.) unless noted otherwise (individual data).

## RESULTS

### Behavior: cocaine self-administration

An initial study on acute effects indicated that VU0364572 produced a moderate reallocation of behavior away from cocaine self-administration towards food-taking across a range of doses, with lower doses increasing food intake while larger doses produced mixed effects that suggested some suppression of food-taking early in the session (see Fig. S2). Acutely, VU0364572 only shifted behavior for the two lower cocaine doses, and did not result in overall reduction in cocaine intake over the session (Table S1). We noticed that cocaine choice remained decreased for a day or two after administration of the low-intermediate doses of VU0364572. More strikingly, many rats that received 1.0 mg/kg VU0364572 subsequently failed to maintain cocaine-taking behavior at their baseline levels. We then explored this phenomenon systematically by recording cocaine vs. food choice behavior for up to four weeks after acute administration of saline or 0.1 or 1 mg/kg VU0364572.

Fig. 2 shows choice behavior between cocaine injections and liquid food reinforcer at baseline and four weeks after acute administration of saline (Fig. 2A-C) or 1 mg/kg VU0364572 (Fig. 2D-F). Rats receiving vehicle showed stable allocation of choice between cocaine and food over the observation period. The effect of cocaine dose was always highly significant (*p*<0.0001) and will not be reported for individual analyses, for brevity. Comparing percent cocaine choice at baseline, on the treatment day, and 7, 14, 21 and 28 days after VU0364572 administration, two-way ANOVA confirmed a significant effect of time after treatment [F(5,186)=8.83, *p*<0.0001] with a cocaine by time interaction [F(20,186)=1.85, *p*=0.02]. Allocation of choice to cocaine-taking was significantly decreased in the VU0364572 treated animals on the treatment day (*p*=0.01), and one (*p*=0.02), two (*p*=0.006), three (*p*=0.0004) and four weeks (*p*<0.0001) later, relative to baseline (Fig. 2D, only baseline and four-week data are shown to avoid crowding). The change in percent choice reflected both a gradual decrease in cocaine injections taken ([F(5,192)=6.03, *p*<0.0001]; Fig. 2E) and an increase in food reinforcers taken ([F(5,192)=6.88, *p*<0.0001]; Fig. 2F). Decreases in cocaine-taking reached significance starting one week after treatment (*p*<0.0004), the increase in food-taking, from two weeks after treatment (*p*<0.005).

**Figure 2.**
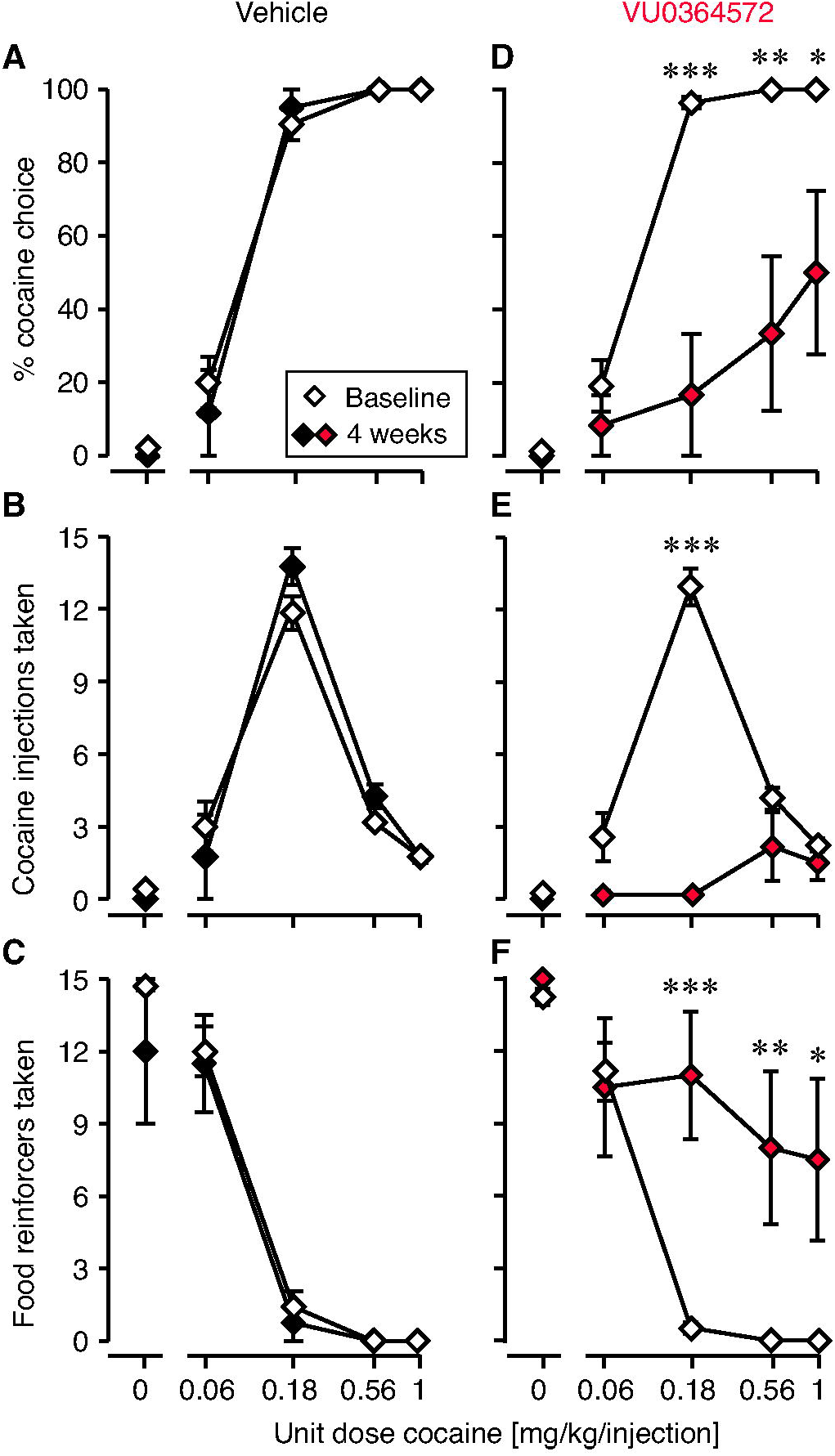
Cocaine self-administration behavior was markedly reduced four weeks after acute VU0364572 (1 mg/kg) administration. In vehicle-treated rats (n=4), cocaine vs. food choice allocation (A), cocaine self-administration (B), and food taking (C) remained constant relative to baseline. In VU0364572-treated rats (n=6-9), cocaine vs. food choice allocation (D) and cocaine self-administration (E) were significantly decreased, and behavior was reallocated to food taking (F) at four weeks post-treatment relative to baseline. \**p*<0.05, \*\**p*<0.01, \*\*\**p*<0.001 vs. baseline.

Also measured as session-wide total allocation of behavior to cocaine vs. food taking and as total intake, after administration of 1 mg/kg VU0364572 rats showed a gradual decrease in cocaine choice [F(6,40)=3.08, *p*=0.01] and cocaine intake [F(6,40)=3.25, *p*=0.01] as a function of time, while food intake increased [F(6,40)=2.83, *p*=0.02]; see Fig. 3A-C for post-hoc significance levels. After administration of vehicle, behavior remained stable over four weeks (no significant effect of time; Fig. 3A-C). Fig. 3D-I shows changes in behavior from baseline to day 28 (four weeks) in individual rats: five of six VU0364572-treated rats showed marked decreases in cocaine choice and intake – and half completely stopped taking cocaine – while saline-treated rats maintained or increased cocaine choice and intake.

**Figure 3.**
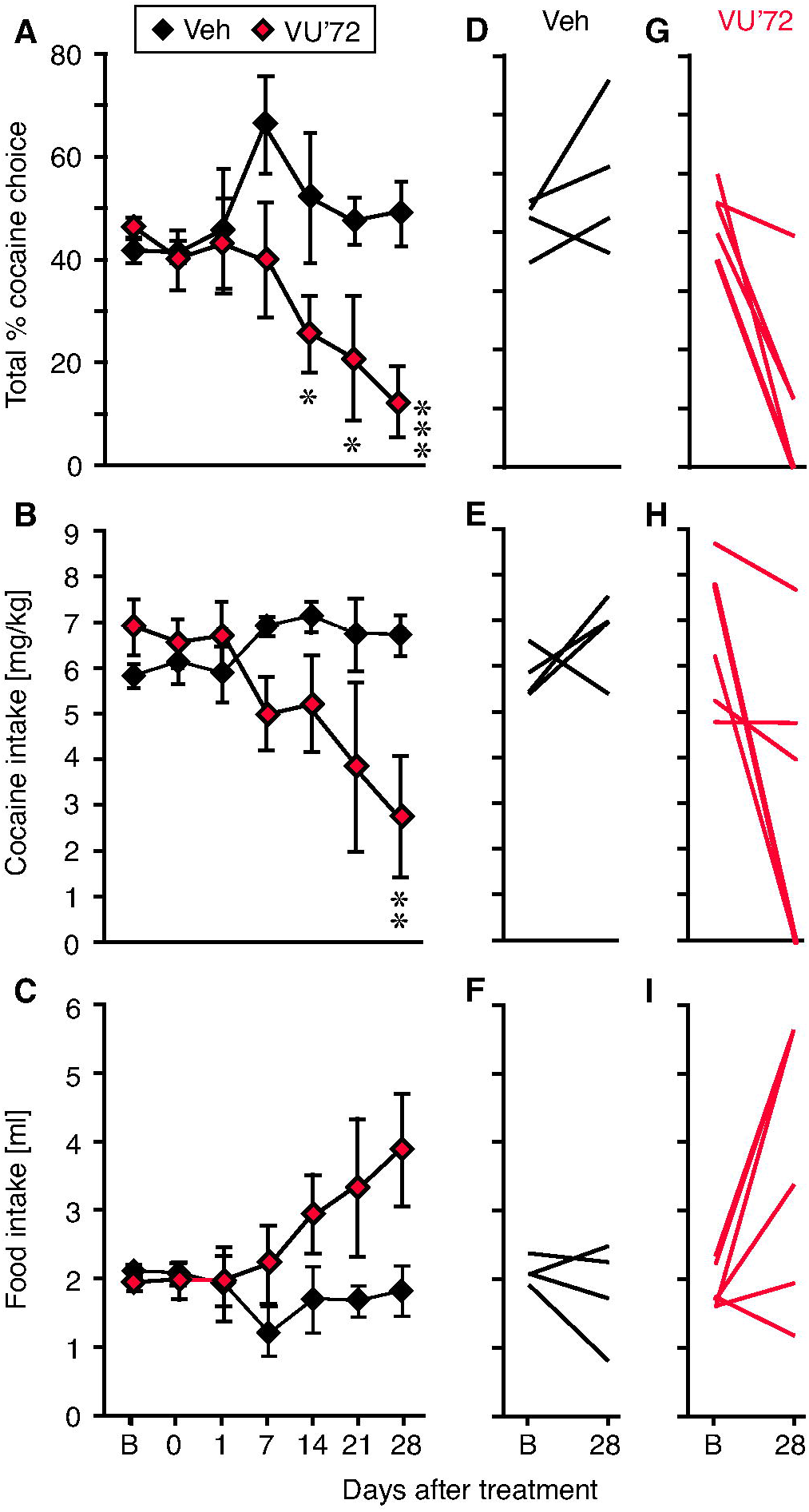
Behavior was gradually reallocated from cocaine taking towards the food reinforcer. Session-wide cocaine vs. food choice allocation (A) and cocaine intake (B) were not significantly decreased on the VU0364572 treatment day, but started decreasing one or two weeks after treatment, while food intake (C) increased. In vehicle-treated rats, behavior remained stable. Changes from baseline to day 28 post-treatment in individual subjects are shown for cocaine choice (D,G), cocaine intake (E,H), and food intake (F,I) in the vehicle group (D-F) and VU0364572 groups (G-I). B: baseline, \**p*<0.05, \*\**p*<0.01, \*\*\**p*<0.001 vs. baseline. Group sizes: saline (Veh), n=4; VU0364572 (VU’72), n=9 baseline to 7 days, n=8 at 14 days, then n=6.

A lower dose of 0.1 mg/kg VU0364572 caused a smaller, more transient decrease in cocaine-taking behavior, peaking 1-2 days after treatment and returning to baseline levels within one or two weeks (Supplemental Fig. S3).

### Microdialysis

Based on evidence that M_1_ receptor stimulation modulates the integration of corticostriatal and dopaminergic signaling in NAc, we then asked if VU0364572 decreased cocaine reinforcement by blunting NAc dopamine release and/or by altering corticostriatal glutamatergic signaling. Rats were treated with 1 mg/kg VU0364572 or vehicle in combination with saline or cocaine, and extracellular levels of dopamine and glutamate were measured four weeks later in vivo by dual-probe microdialysis in mPFC and NAc, after acute administration of 15 mg/kg cocaine. VU0364572 was never administered on the test day.

Extracellular dopamine levels were related to cocaine history and VU0364572 history both in the mPFC (cocaine [F(1,20]=10.1, *p*=0.005], VU0364572 [F(1,20]=6.64, *p*=0.02], interaction [F(1,20]=26.1, *p*=0.0001]) and in the NAc (cocaine [F(1,20]=9.66, *p*=0.006], VU0364572 [F(1,20]=43.2, *p*<0.0001], interaction [F(1,20]=6.23, *p*=0.02]). In the mPFC, acute cocaine administration increased dopamine levels above baseline in the cocaine-experienced (sensitized, Fig. 4B) vehicle-treated animals [F(7,35)=11.0, p<0.0001] but not in cocaine-naïve animals (Fig. 4A). The dopamine increase in the cocaine-experienced animals was completely blocked by VU0364572 pre-treatment (effect of pretreatment [F(1,10]=18.3, *p*=0.002]; Fig. 4B). Fig. 4C shows overall %increase in dopamine over baseline, which was related to VU0364572 history [F(1,20]=6.25; *p*=0.02] and cocaine history [F(1,20]=8.66; *p*=0.008] with significant interaction [F(1,20]=11.6; *p*=0.003). Vehicle-treated cocaine-experienced rats had higher cocaine-stimulated mPFC dopamine levels relative to vehicle-treated naïve rats (*p*=0.0006) and relative to VU0364572-treated cocaine-experienced rats (*p*=0.001).

**Figure 4.**
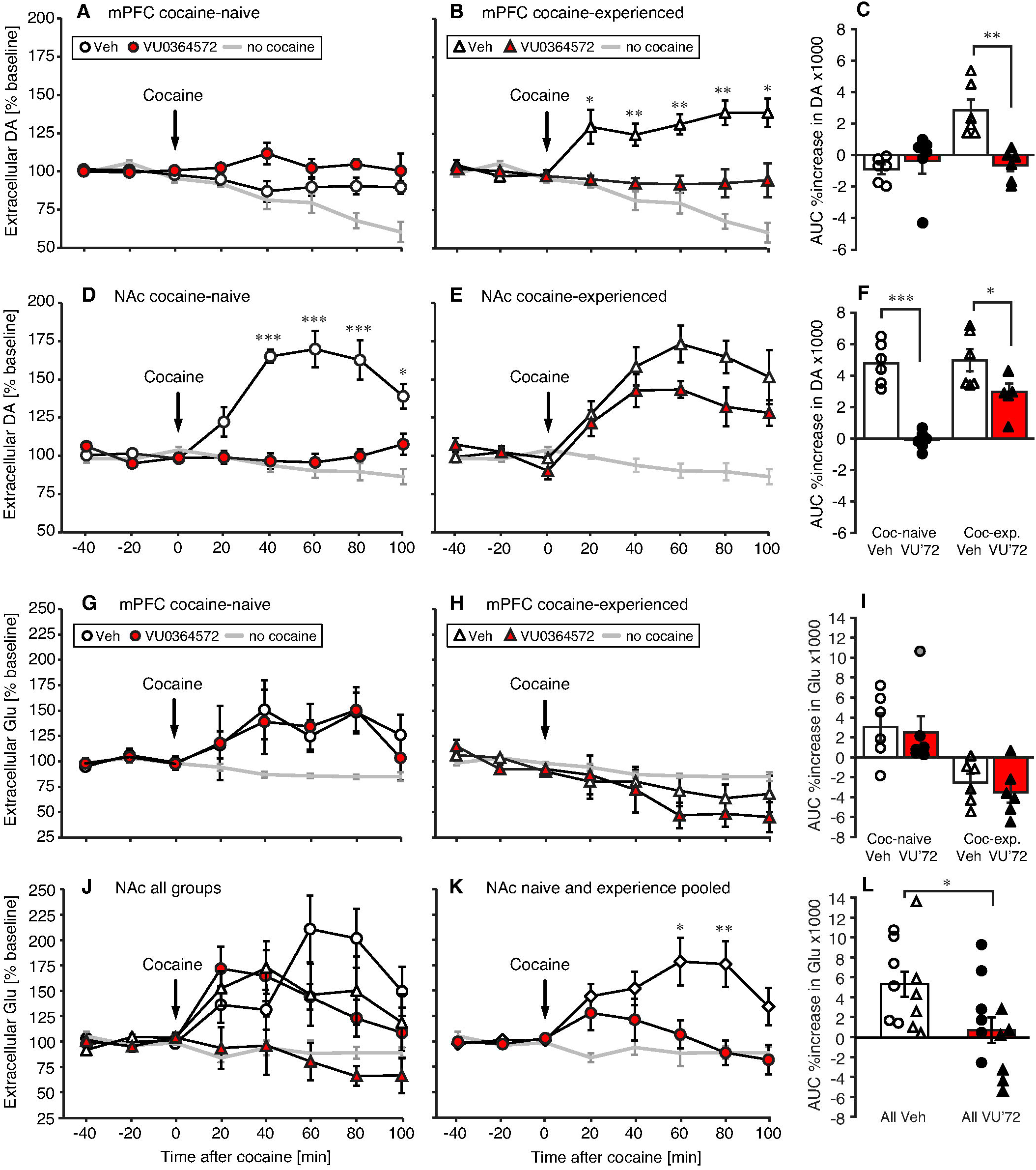
Rats treated with VU0364572 (VU’72) four weeks earlier showed blunted dopamine responses to cocaine in PFC (A-C) and NAc (D-F). Acute cocaine administration (15 mg/kg, arrow) caused increased dopamine levels in NAc in both cocaine-naïve and cocaine-experienced (D,E) rats, measured by in vivo dual probe microdialysis. The VU0364572 treatment also blunted glutamate (Glu) responses to cocaine in NAc (J-L). Acute cocaine caused increased glutamate levels in NAc in cocaine-naïve and cocaine-experienced (J) rats. Overall increases in dopamine (C,F) and glutamate (I,L) levels as area under the curve of 0-100 min after cocaine injection; points represent individual rats. n=6, \**p*<0.05, \*\**p*<0.01, \*\*\**p*<0.001 vs. vehicle (Veh).

In the NAc, cocaine administration increased dopamine levels above baseline in both cocaine-naïve [F(7,35)=17.7, p<0.0001] and cocaine-experienced animals [F(7,35)=10.7, p<0.0001] treated with vehicle (Fig. 4D,E). VU0364572 treatment significantly blunted the cocaine-stimulated increase in extracellular NAc dopamine in both cocaine-naïve [F(1,10]=62.9, *p*<0.0001] and cocaine-experienced [F(1,10]=6.13, *p*=0.03] rats, but with different efficacy. In the cocaine-naïve animals, prior VU0364572 treatment completely blocked the dopamine increase, while the cocaine-experienced animals still showed a significant dopamine increase [F(7,35)=6.98, p<0.0001]. Fig. 4F shows overall %increase in dopamine over baseline, which was related to VU0364572 treatment [F(1,20]=41.4; *p*<0.0001] and cocaine history [F(1,20]=9.54; *p*=0.006] with interaction [F(1,20]=7.24; *p*=0.01]. Cocaine-stimulated NAc dopamine levels were decreased in VU0364572-treated rats relative to vehicle-treated rats in both cocaine-naïve (*p*<0.0001) and cocaine-experienced (*p*=0.049) animals.

Extracellular glutamate levels in the mPFC were significantly related to cocaine history [F(1,20)=23.5, p=0.0001], although the effect of acute cocaine administration did not reach statistical significance relative to baseline levels in either group (Fig. 4G,H). There was no significant effect of VU0364572 history or interaction. There was a statistical outlier (over two standard deviations above the mean) in the naïve/VU0364572 group, depicted with a grey point in Fig. 4I, but analyzing the data without this outlier did not reveal a significant effects of VU0364572 history. Fig. 4I shows overall %increase in glutamate over baseline, which was again related to only cocaine [F(1,20)=20.9; *p*=0.0002]. Cocaine-experienced rats had significantly lower cocaine-stimulated mPFC glutamate levels than cocaine-naïve rats whether they were treated with saline (*p*=0.007) or VU0364572 (*p*=0.01).

Acute cocaine administration increased NAc glutamate levels in vehicle- and VU0364572-treated cocaine-naïve rats ([F(7,35)=5.91, p=0.0001] and [F(7,35)=4.07, p=0.002], respectively) and in vehicle-treated cocaine-experienced rats [F(7,35)=2.59, p=0.03], but the increase was completely abolished in the VU0364572-treated cocaine-experienced rats (Fig. 4J). Extracellular glutamate levels in the NAc were lower in rats previously treated with VU0364572 [F(1,20]=7.39, *p*=0.01], while effects of cocaine history and interactions were not significant (Fig. 4J). Data were therefore analyzed post-hoc with cocaine-naïve and cocaine-experienced animals pooled (see Fig. 4K for significant time-points). Fig. 4L shows overall %increase in glutamate over baseline, which was related to only VU0364572 treatment [F(1,20)=7.37; *p*=0.01]. VU0364572-treated rats had lower cocaine-stimulated NAc glutamate levels, naïve and experienced rats combined (p=0.02).

### Dopamine uptake

To determine whether the large effects of VU0364572 treatment on extracellular dopamine levels were due to altered dopamine transporter function, we assessed dopamine uptake in striatal tissue from rats that received the same treatments as in the microdialysis study. Dopamine uptake was related to dopamine concentration [F(5,60)=59.2, p<0.0001] with no significant effect of cocaine or VU0364572 history (Fig. 5). V_max_ and K_M_ did not differ significantly between groups. There was a non-significant apparent decrease in dopamine uptake in the cocaine-experienced vehicle-treated rats relative to cocaine-naïve rats, consistent with previous investigations. The VU0364572-treated cocaine-experienced rats showed dopamine uptake indistinguishable from naïve rats. Because of this possible effect of VU0364572 treatment in the cocaine-experienced group, the experiment was subsequently repeated in cocaine-experienced animals only (n=4, data not shown). This showed the same trend, and overall, Vmax was 39% higher in VU0364572-treated cocaine-experienced rats than in vehicle controls (p=0.08), with no effect of VU0364572 treatment in cocaine-naïve rats (Vmax 101.8% of controls, p>0.9).

**Figure 5.**
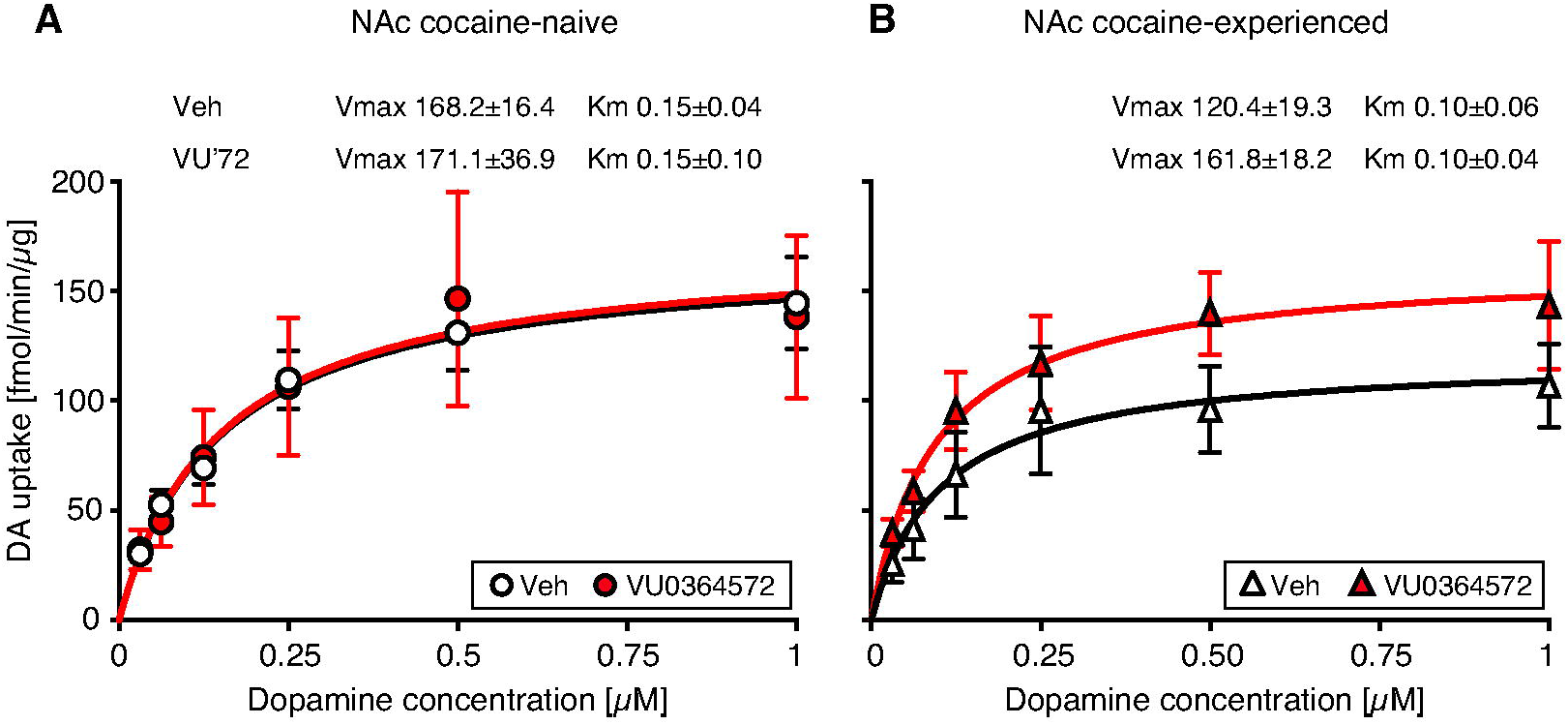
Saturation curve for dopamine uptake in synaptosomes from accumbens of (A) cocaine-naïve and (B) cocaine-experienced rats previously treated with vehicle (Veh) or VU0364572 (n=4). Best-fit values ± s.e.m. of the maximal dopamine uptake capacity V_max_ in fmol/min/μg and apparent affinity K_M_ in μM are indicated for each group.

## DISCUSSION

We discovered large effects on cocaine self-administration behavior and cocaine-induced dopamine and glutamate outflow weeks after treatment with the M_1_ agonist VU0364572. While much remains to be elucidated about the mechanism underlying these unusual delayed effects, our findings are consistent with M_1_-dependent modulation in the addiction pathway that comprises the glutamatergic corticostriatal projections.

### Cocaine self-administration behavior

Acutely, VU0364572 produced small decreases in cocaine self-administration consistent with functional antagonism of the reinforcing effects of cocaine: VU0364572 decreased self-administration of low to moderate doses of cocaine but not high doses, with no effect on overall intake levels. This is similar to the effects of the M_1_/M_4_-preferring agonist xanomeline in the same assay [23] and consistent with attenuation of cocaine discrimination by M_1_ and M_4_ stimulation [22,33]. In the course of the experiment, it became apparent that VU0364572 could suppress cocaine self-administration in a much more pronounced manner, days to weeks after dosing. At a low dose, VU0364572 produced moderate decreases in cocaine-taking for 1-2 days after treatment, reminiscent of xanomeline [23]. These effects were magnified and prolonged with a higher VU0364572 dose, which over four weeks caused a remarkable decrease in cocaine-taking behavior – half the rats stopped taking cocaine entirely. We observed no adverse effects and rats continued to press the lever reinforced with food, indicating a lack of motor impairment or stimulus control disruption. Importantly, the delayed effects differed qualitatively from acute effects by showing a reduction in cocaine intake with no evidence of increased intake at the higher cocaine doses. Rather, the delayed effect is similar to the effect of removing the cocaine syringe in this assay [34], consistent with VU0364572 gradually reducing (removing) cocaine’s reinforcing effects. Increased reinforcing effect of the food is unlikely to account for the effects, since we never observed increased food-reinforced operant behavior after M_1_ receptor stimulation [22,26,33], and others reported increased food-reinforced behavior after *blocking* M_1_ receptors [35].

### Microdialysis, dopamine uptake, and suggested mechanism

Vehicle-treated control rats yielded findings consistent with previous reports: cocaine significantly increased mPFC dopamine in cocaine-experienced rats only, increased NAc dopamine and glutamate in both cocaine-naïve and -experienced rats, and failed to consistently increase mPFC glutamate [36–40]. VU0364572 treatment four weeks before testing completely prevented cocaine-induced increases in extracellular mPFC dopamine and NAc glutamate in cocaine-experienced rats. Both effects are consistent with suppressed cocaine self-administration: several studies indicate that PFC dopamine, but not glutamate, plays an important role in cocaine seeking and taking in cocaine-experienced rats, whereas in NAc, glutamate, but not dopamine signaling, appears to mediate these behaviors [36,41–46]. The VTA is the main source of dopamine in both mPFC and NAc [47], but VU0364572 did not affect dopamine response equally in both regions, perhaps making modulation in VTA dopamine neurons less likely. Further, M_1_ receptors are not expressed on VTA dopamine neurons [16,48]. Taken together, we propose that M_1_-mediated modulation of corticostriatal glutamatergic pathways best explains our findings. Acutely, M_1_ receptor stimulation was shown to postsynaptically increase NMDA but not AMPA receptor-mediated responses in MSN, and to modulate PFC and corticostriatal neuroplasticity [28,49–53]. A balancing of relative AMPA/NMDA transmission, which is shifted following cocaine exposure [2,45,54], could be a way in which M_1_ stimulation opposed cocaine’s effect, although it is unclear how those acute M_1_ receptor-dependent actions and our delayed effects are related.

Cocaine-naïve, VU0364572-treated rats were “immune” to cocaine-induced NAc dopamine outflow. The fact that VU0364572 also blunted the cocaine-induced dopamine response in cocaine-naïve rats indicates that at least some effects reflect modulation of the circuitry independent of cocaine-induced plasticity. These results extend our finding that M_1_/M_4_ receptor stimulation facilitated extinction of cocaine seeking also when not paired with cocaine exposure or extinction learning [26], although in those studies all animals were cocaine-experienced. It would be interesting to test whether M_1_ stimulation reduces subsequent acquisition of cocaine self-administration in drug-naïve animals.

The large reductions in cocaine-induced dopamine outflow in VU0364572-treated rats cannot be attributed to up-regulation of cell-surface DAT, as VU0364572 treatment had no effect on dopamine uptake in cocaine-naïve rats, and only a non-significant trend in cocaine-experienced rats. Our results in the vehicle-treated rats are consistent with preclinical [55] and clinical [56] studies showing modest decreases in DAT availability weeks after cocaine self-administration or passive exposure, an effect that is variable in humans and likely depends on the duration of abstinence [57–60]. While inconclusive, VU0364572 treatment may have prevented/attenuated this cocaine-induced change in DAT function; testing at intervals when cocaine-induced changes in DAT availability are more robust is needed to answer this question.

### Limitations and future directions

Due to probe size, our microdialysis data cannot distinguish between effects in the prelimbic vs. infralimbic mPFC, and between the NAc core, NAc shell, or medioventral caudate putamen, although probe tips were located in the NAc core. Because prelimbic mPFC to NAc core projections and infralimbic mPFC to NAc shell projections are thought to modulate cocaine seeking differentially [2,3], it would be interesting to extend our studies with more spatially focused methodologies. There are differences between effects of self-administered and experimenter-administered cocaine, and the rats self-administering cocaine did not have a 4-week forced abstinence period. This precludes a direct comparison between the microdialysis and uptake experiments and the behavioral experiments, limiting what conclusions can be drawn about mechanism. Conversely, the demonstration that M_1_ receptor stimulation opposed the effects of cocaine regardless of prolonged abstinence, extinction training, or mode and route of cocaine administration strengthens our confidence in the robustness of the effects and the potential clinical utility of the approach.

We observed no adverse effects of VU0364572 at the doses tested. Excessive M_1_ stimulation may cause adverse effects including nausea, cognitive impairment, or pro-convulsant effects, which could be worrisome clinically especially when combined with cocaine use [61–66]. However, adverse effects may not be a great concern given that we obtained pronounced effects at relatively low doses of VU0364572, which was tolerated in rats at 10, 32, and even 56 mg/kg [28,67,68]. M_1_ receptor stimulation has shown promise improving cognitive performance in rodents and, in that context, it was suggested that M_1_ positive allosteric modulators (PAMs) may retain efficacy while lacking cholinergic toxicity [64,69]. Whether pure M_1_-PAMs can replicate the current effects of the bitopic (allosteric and orthosteric) PAM/agonist VU0364572 [27] is worth exploring. It is possible that VU0364572 has a particularly useful pharmacological profile, as allosteric M_1_ agonists differ in activation of specific signaling pathways [65,67,70]. Nevertheless, we have observed acute “anti-cocaine” effects using several structurally distinct M_1_ agonists/PAMs, suggesting a common M_1_-dependent action [23,26,33].

Finally, corticostriatal glutamatergic transmission is also modulated (generally inhibited) by stimulation of presynaptic muscarinic M_4_ receptors in NAc [71,72], and we have observed additive or synergistic effects on cocaine-induced behaviors by stimulating both M_1_ and M_4_ receptors, relative to either alone [22,26,33]. Thus, a combination of highly selective M_1_ and M_4_ receptor agonists or PAMs may show additional benefits in modulating addiction-related effects of cocaine.

## Conclusions

These tardive effects of the muscarinic M_1_ receptor agonist may seem unusual in the context of typical preclinical pharmacology findings. However, they are in line with the clinically observed therapeutic effects of some types of medications in psychiatry, such as antidepressant and antipsychotic drugs. Viewed in that perspective, the authors suggest that any medication capable of ameliorating neurological maladaptations incurred in the development of a drug addiction would perhaps be unlikely to show its full effect acutely. Conversely, “classic” short-acting drugs require excellent adherence to treatment over time to be effective at minimizing relapse. The present findings not only reinforce our interest in M_1_ receptor agonists/PAMs as potential pharmacotherapies in cocaine addiction (and perhaps other addictions), but also suggest that it may be time to revisit the typical acute-treatment approach to evaluating new targets, focusing instead on longer windows of evaluation.

## AUTHOR CONTRIBUTIONS

MT conceptualized the studies. MT, PW, and KLJ designed the experiments, analyzed the data, and interpreted the data. PW conducted the microdialysis studies, KLJ conducted the dopamine uptake experiments. MT directed the self-administration experiments. MT wrote the manuscript with assistance from KLJ and PW.

## ACKNOWLEDGMENTS

We thank professors P. Jeffery Conn and Craig W. Lindsley (Vanderbilt University) for the gift of VU0364572 for these studies. We thank Christopher Adam, Benoît Niclou, Kevin Stoll (bevahior), and Saiy Kiasari, Dr. Gitta Wörtwein, Anne-Marie Paulsen (microdialysis) for technical assistance.

