## Supplemental Figure 1 for "Effects of muscarinic M_1_ receptor stimulation on reinforcing and neurochemical effects of cocaine in rats"

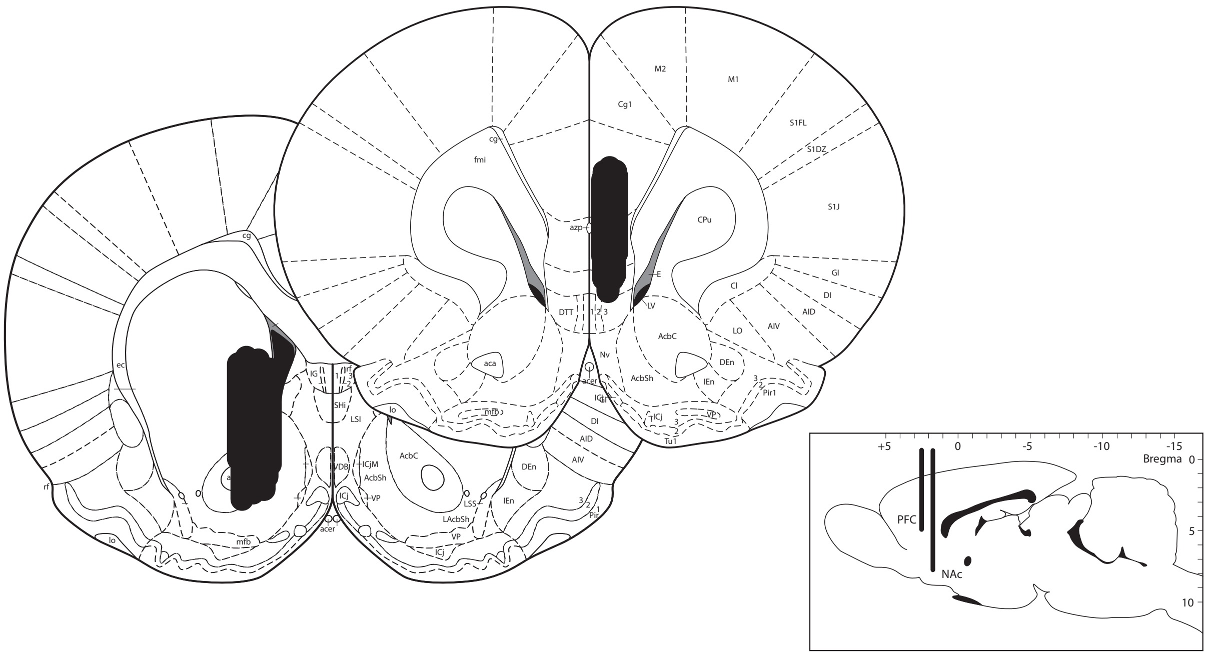

**Supplemental Figure S1**
Individual probe placements for dual-probe microdialysis in mPFC (shown in front) and NAc (shown in back). Insert shows target coordinates in a sagittal view. Atlas from Paxinos and Watson 2005.

Paxinos, G. and Watson, C. (2005). The Rat Brain in Stereotaxic Coordinates. *Compact 6th Edition, Academic Press, New York*, 400.
