## Supplemental Figure 2 for "Effects of muscarinic M_1_ receptor stimulation on reinforcing and neurochemical effects of cocaine in rats"

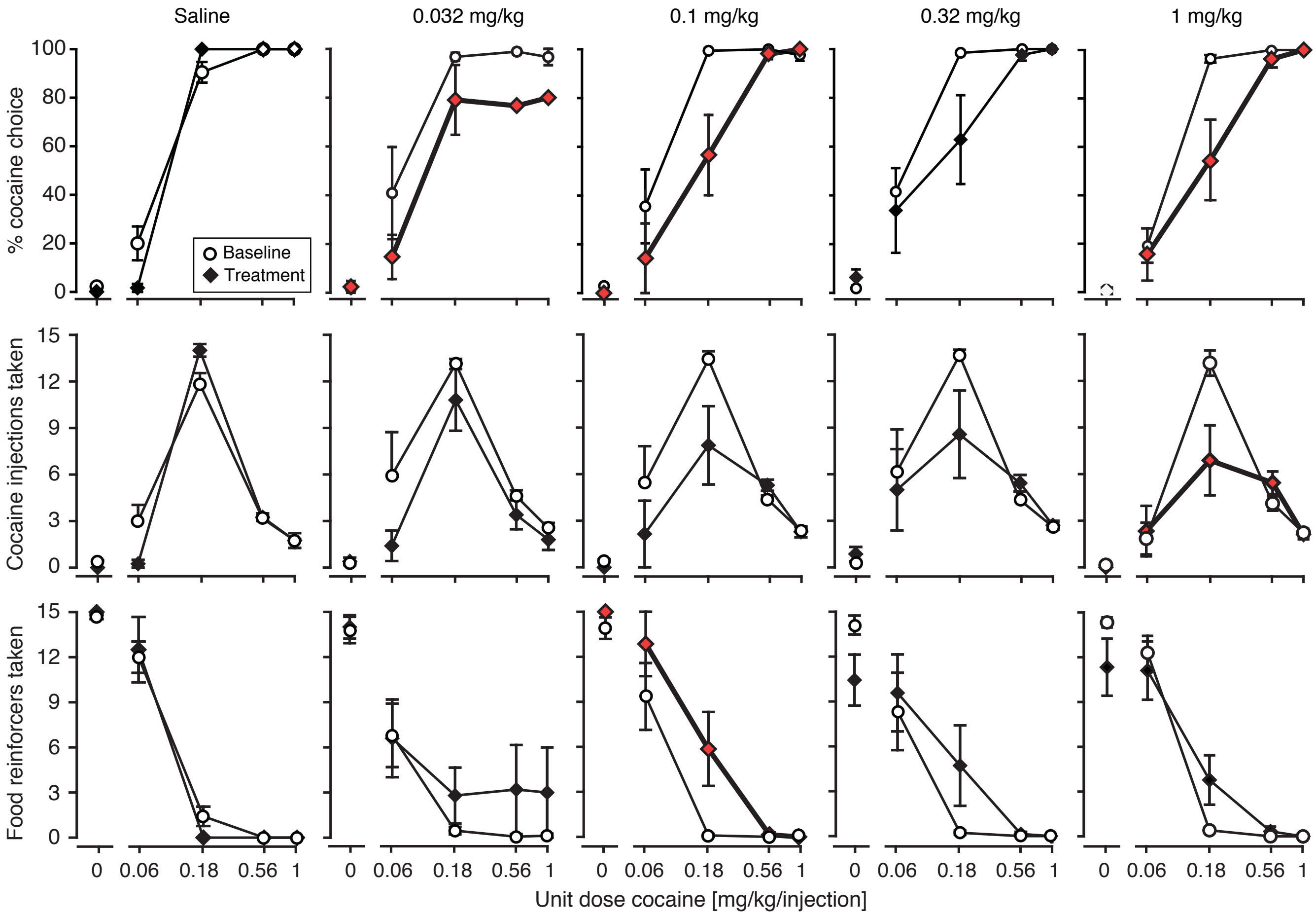


**Supplemental Fig. S2**

All VU0364572 doses tested produced a small apparent reallocation of behavior away from cocaine self-administration towards food taking. The shift in choice allocation reached significance at 0.032 mg/kg [F(1,36)=5.24, *p*=0.03], 0.1 mg/kg [F(1,45)=4.72, *p*=0.04], and 1 mg/kg [F(1,72)=5.58, *p*=0.02], the latter with a significant cocaine dose by treatment interaction [F(1,72)=3.71, *p*=0.008].

Effects on cocaine injections taken only showed a significant cocaine dose by treatment interaction at the highest dose, 1 mg/kg [F(1,72)=3.88, *p*=0.007]. The interaction and lack of main effect of treatment reflect a shift in cocaine taking from lower doses towards higher doses of cocaine, consistent with functional antagonism of cocaine reinforcement. However, the relatively short half-life of VU0364572 (46 min in rats; Lebois et al. 2011) may also contribute to the lack of effect at higher cocaine doses: VU0364572 levels would have been lower towards the end of the 108-min session.

Effects on food appeared biphasic, with a non-significant increase at the lowest dose, reaching a significant increase at 0.1 mg/kg [F(1,45)=5.62, *p*=0.02]; then mixed decrease/increase at the higher doses, with an initial suppression of food-taking early in the session followed by an increase in later components.

Significant effects are indicated by red symbols and bold lines; no comparisons reached significance post-hoc for individual cocaine doses. Group sizes: from left to right n=4, n=5, n=7, n=7, n=9. Saline administration had no significant effect on percent choice allocation, cocaine injections taken, or food reinforcers taken.

Lebois EP, Digby GJ, Sheffler DJ, Melancon BJ, Tarr JC, Cho HP, et al. (2011): Development of a highly selective, orally bioavailable and CNS penetrant M1 agonist derived from the MLPCN probe ML071. *Bioorg Med Chem Lett.* 21:6451-6455.
