## Supplemental Figure 3 for "Effects of muscarinic M_1_ receptor stimulation on reinforcing and neurochemical effects of cocaine in rats"

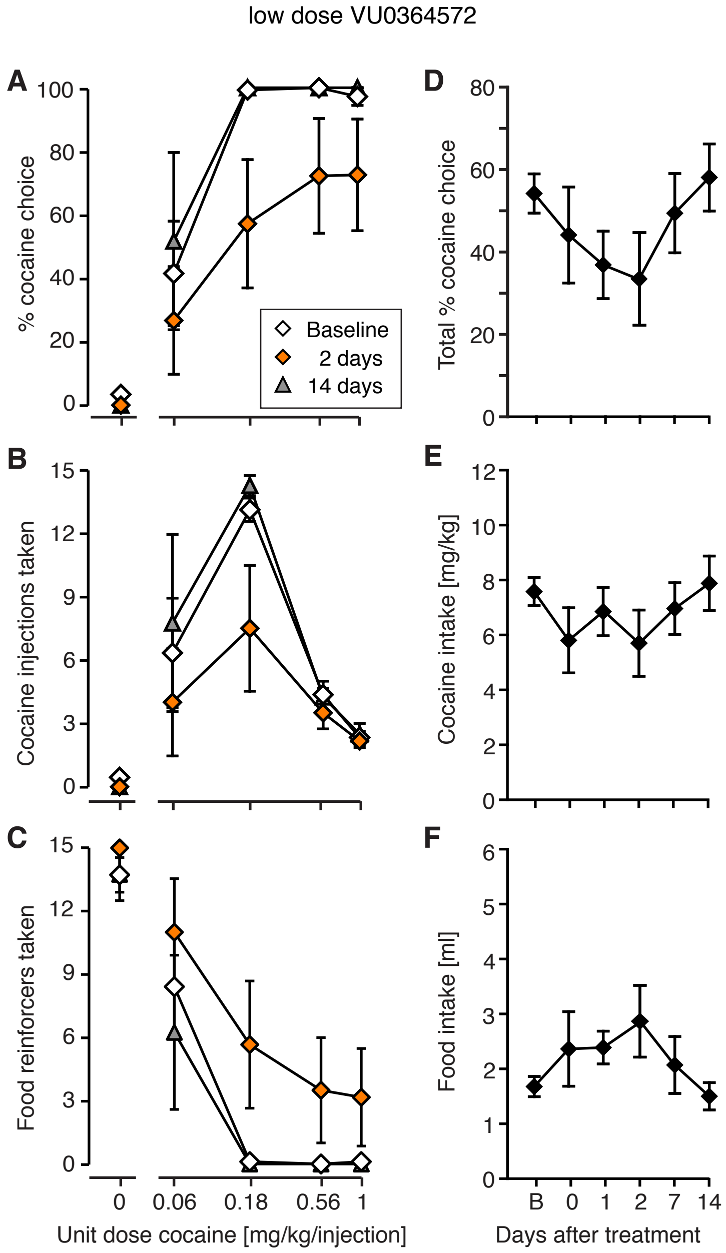


**Supplemental Figure S3**

Cocaine self-administration behavior was transiently reduced after administration of a lower dose of VU0364572, 0.1 mg/kg (n=7). This was reflected by a significant effect of time on %cocaine choice [$F(3,95)=2.98$, $p=0.4$] over the first two days after VU364572 administration. On day 2 post-treatment (orange diamonds), cocaine choice allocation (A, [$F(1,45)=10.1$, $p=0.003$]) and cocaine injections taken (B, [$F(1,45)=4.61$, $p=0.04$]) were reduced and food reinforcers taken were increased (C, [$F(1,45)=9.06$, $p=0.004$]), relative to baseline (open diamonds). After two weeks, behavior had returned to baseline levels (A-C, grey triangles, n=5). Some rats were tested for a full four weeks post-treatment, and did not show further suppression of cocaine taking (n=3, data not shown).

Reallocation of cocaine-taking behavior towards food-taking behavior peaked 1-2 days after treatment, and returned to baseline after 1-2 weeks (D-F). However, analysis of session-wide choice allocation (D), cocaine intake (E), and food intake (F) over two weeks did not reach statistical significance. B: baseline.
