## Supplemental methods for "Effects of muscarinic M_1_ receptor stimulation on reinforcing and neurochemical effects of cocaine in rats"

### Microdialysis

Relative glutamate concentrations were detected by a C18 reversed-phase column 3.5μm Zorbax 300SB C18 analytical column (4.6mm × 150mm id, Agilent Technology Inc., USA) and gradient fluorescence HPLC detection. The dialysate was pre-column derivatized with *o*-phthalaldehyde (Merck, Darmstadt, Germany). Mobile phase A consisted of 70 mM $Na_2HPO_4$ in 50 μM of EDTA 0.15%(v/v) tetrahydrofurane (Sigma-Aldrich) and 20% (v/v) methanol (Sigma-Aldrich); pH was adjusted to 7.2 with 0.1 M acetic acid. Mobile phase B was 100% methanol. The gradient elution was performed at room temperature using the following conditions: 0 min 15% B (flow rate of pump B: 0.65 mL/min), 2.00 min 22% B (flow rate of B: 0.65 mL/min) for A+B after 15 min 10% B mobile phase. The excitation and emission wavelengths were 350 and 450 nm, respectively. Known standards were used to identify and quantify the glutamate peaks in the HPLC chromatogram. The HPLC system consisted of a Shimadzu chromatograph (LC-20AD, Tokyo, Japan) equipped with an autosampler, a degasser (DGU-20A3), a gradient pump (LC-20AD), a fluorescence detector model RF-20Axs. Characteristic retention times for glutamate were 4.9 min. The peak areas of glutamate were integrated with the calibration curve of known standards.

### Data analysis

*Behavior.* Numbers of cocaine injections and food reinforcers taken and cocaine injections as percent choice over total reinforcers were recorded for each component, and were analyzed for each VU0364572 dose by two-way ANOVA with cocaine dose and time after treatment as repeated-measures factors. Significant effects of time or cocaine by time interactions were followed by simple effects analysis on each time point vs. baseline, by two-way ANOVA. Session-wide cocaine intake (mg/kg), food intake (ml), and total percent cocaine choices were calculated and were analyzed for each VU0364572 dose by one-way ANOVA with time after treatment as repeated-measures factor. Significance at individual cocaine doses (two-way analyses) or time points (one-way analyses) was determined by two-tailed paired-sample t-test corrected for false discovery rate (Benjamini-Hochberg, 5% false discovery rate limit).

*Microdialysis*. The effect of acute cocaine administration on the test day was evaluated as a pre-planned analysis by one-way repeated measures ANOVA in each treatment group and for each brain region and neurotransmitter, and was indicated by a significant change in neurotransmitter level as a function of time. Effects of treatment history were evaluated for each brain region and neurotransmitter separately, by three-way ANOVA with cocaine and VU0364572 as between-subjects factors and time after cocaine injection (0-100min) as repeated-measures factor. Significant effects of cocaine history, VU0364572 history, or their interaction, were further evaluated by two-way ANOVA with cocaine or VU0364572 history and time as factors, followed by false discovery rate-corrected unpaired-sample t-test. In addition, area under the curve (AUC) for increase in neurotransmitter levels above baseline 0-100 min after cocaine administration was calculated (trapezoid method) and analyzed by two-way ANOVA with cocaine and VU0364572 history as factors, followed by false discovery rate-corrected unpaired-sample t-test.

*Dopamine uptake*. Uptake was analyzed by three-way ANOVA with cocaine and VU0364572 as between-subjects factor and dopamine concentration as a repeated-measures factor. The fitted values for $V_{max}$ and $K_M$ were compared by two-way ANOVA with cocaine and VU0364572 as factors.

Level of significance was set at $\alpha$=0.05. Data are presented as group means ± standard error of the mean (s.e.m.) unless noted otherwise (individual data). Power analyses were performed using G*power 2 (Faul et al. 2007). Behavior: a priori analysis based on effects of xanomeline in the choice assay (Thomsen et al. 2014) yielded a required n=4-6. Microdialysis: post-hoc analysis yielded a power of >0.99 for all four analyses, dopamine and glutamate in mPFC and NAc. Dopamine uptake: a priori analysis based on our previous studies in mice yielded a minimum n=3; however variation was larger than expected. Post-hoc power analysis showed low power in the cocaine-experienced groups, we therefore repeated the experiment in a second cohort, cocaine-experienced only.

Faul F, Erdfelder E, Lang AG, Buchner A (2007): G*Power 3: a flexible statistical power analysis program for the social, behavioral, and biomedical sciences. *Behav Res Methods*. 39:175-191.

Thomsen M, Fulton BS, Caine SB (2014): Acute and chronic effects of the M1/M4-preferring muscarinic agonist xanomeline on cocaine vs. food choice in rats. *Psychopharmacology (Berl)*. 231:469-479.
