## Supplemental Table 1 for "Effects of muscarinic M_1_ receptor stimulation on reinforcing and neurochemical effects of cocaine in rats"

**Supplemental Table S1**

Total cocaine and food reinforcer intake per session after acute treatment with VU0364572

| Dose VU0364572 | Cocaine intake [mg/kg/session] | Food reinforcer intake [ml/session] | Total cocaine choice [% choices/session] |
|---|---|---|---|
| Saline | | | |
| baseline | 5.82±0.26 | 2.11±0.10 | 41.70±2.34 |
| treatment day | 6.10±0.45 | 2.06±0.16 | 41.44±2.16 |
| 0.032 mg/kg | | | |
| baseline | 7.81±0.70 | 1.60±0.16 | 54.83±5.35 |
| treatment day | 5.73±1.46 | 2.22±0.48 | 39.58±7.41 |
| 0.1 mg/kg | | | |
| baseline | 7.58±0.51 | 1.68±0.18 | 54.18±4.76 |
| treatment day | 5.80±1.19 | 2.36±0.68 | 44.11±11.7 |
| 0.32 mg/kg | | | |
| baseline | 7.86±0.63 | 1.68±0.16 | 53.55±4.97 |
| treatment day | 7.58±0.91 | 1.86±0.37 | 46.76±10.1 |
| 1.0 mg/kg | | | |
| baseline | 6.89±0.61 | 1.95±0.11 | 45.96±1.93 |
| treatment day | 6.53±0.54 | 1.99±0.27 | 39.62±5.84 |

No dose of VU0364572 significantly affected cocaine intake (mg/kg/session) or food reinforcer intake (ml/session) on the treatment day, relative to baseline. Values are group means ± s.e.m. Group sizes as in Fig. S1.
